# Predicting protein subcellular location using learned distributed representations from a protein-protein network

**DOI:** 10.1101/768739

**Authors:** Xiaoyong Pan, Lei Chen, Min Liu, Tao Huang, Yu-Dong Cai

## Abstract

Functions of proteins are in general related to their subcellular locations. To identify the functions of a protein, we first need know where this protein is located. Interacting proteins tend to locate in the same subcellular location. Thus, it is imperative to take the protein-protein interactions into account for computational identification of protein subcellular locations.In this study, we present a deep learning-based method, node2loc, to predict protein subcellular location. node2loc first learns distributed representations of proteins in a protein-protein network using node2vec, which acquires representations from unlabeled data for downstream tasks. Then the learned representations are further fed into a recurrent neural network (RNN) to predict subcellular locations. Considering the severe class imbalance of different subcellular locations, Synthetic Minority Over-sampling Technique (SMOTE) is applied to artificially boost subcellular locations with few proteins.We construct a benchmark dataset with 16 subcellular locations and evaluate node2loc on this dataset. node2loc yields a Matthews correlation coefficient (MCC) value of 0.812, which outperforms other baseline methods. The results demonstrate that the learned presentations from a protein-protein network have strong discriminate ability for classifying protein subcellular locations and the RNN is a more powerful classifier than traditional machine learning models. node2loc is freely available at https://github.com/xypan1232/node2loc.

## 1. Introduction

Functions of proteins are in general related to their subcellular locations. The specific location of a protein can provide a guide for the design of effective new drugs, e.g. plasma membrane proteins located in extracellular space are easily accessible by drug molecules. Currently there exist many experimental methods to determine the locations of proteins. However, these experimental methods are costly and time-consuming to determine subcellular locations of all proteins in the post-genomics era. Fortunately, there are many experimentally verified locations of proteins in Swiss-Prot database [1], and they can serve as training data for machine learning models. Thus, computational identification of protein compartments using machine learning is imperative for further functional analysis of proteins and has clinical potentials.

In the past decade, many computational methods have been developed to predict protein subcellular locations from sequences using machine learning [2, 3]. For example, Cai and Chou [4] combine functional domain composition and support vector machine to predict the subcellular locations. Furthermore, they integrate pseudo amino acid composition and sequence-order effect to improve the prediction accuracy [5]. Euk-mPLoc[2] trains a fused classifier to predict Eukaryotic Protein subcellular location by integrating multiple sites. Constructing informative features is very crucial for protein subcellular location prediction. For example, Hum-mPloc 3.0 [6] combines the GO information and functional domain features to enhance prediction performance. To remove some redundant features, Li et al. first use feature selection to select the discriminant features, which are further fed into machine learning classifier to predict subcellular locations [7].

Most of the previous methods only consider the information from the protein itself and ignore the biological context, especially under the context of protein-protein interactions. It is observed that proteins that interact tend to reside within the same subcellular locations [8]. Proteins with the similar functional annotations in a protein-protein network may have a complex relationship, which provide informative clues for subcellular locations. Thus, incorporating the context of the protein-protein interaction (PPI) network into feature engineering can extract the discriminative features for classifying subcellular compartments.

Recently deep learning has been extensively applied in computational biology [9]. For example, DeepBind uses convolutional neural network (CNN) [10] to infer binding preference of proteins [9]. iDeep uses hybrid CNN and deep belief network (DBN) to integrate multiple sources of data to further improve the prediction performance of RNA-protein interaction [11]. iDeepS[12] identifies both binding sequence and structure preference of RNA-binding proteins using a CNN and a long short temporary memory network (LSTM) [13]. iDeepE further improves the prediction performance by combining local and global CNNs [14]. Beside predicting protein-RNA binding sites, CNNs are also used in other biological tasks. DeepSEA trains a CNN to predict noncoding variant effects from sequences [15]. DanQ combines a CNN and a bidirectional LSTM to improve the prediction performance [16].

However, CNNs and RNNs cannot handle structured data, like network data. Thus, some network embedding methods have been developed to extract high-level features from the network [17, 18]. For example, node2vec uses shallow and linear techniques in word2vec [19] to capture complex relationship and preserve the network structure [17]. Embedding of proteins projects proteins into a lower dimensional space, in which each protein is represented by a vector instead of one-hot encoding or other hand-designed features.

In addition, there exist data imbalances in different subcellular locations as shown in Table 1, which may make the model be preferred to majority classes. How to reduce the impact of imbalanced class on model performance is also very important. Many over-sampling techniques, such as Synthetic Minority Over-sampling Technique (SMOTE)[20] or Supervised Over-Sampling [21], have been used to reduce the impact of imbalanced classes. SMOTE synthetizes the same number of samples for the existing minority classes to those majority class. In this study, we applied SMOTE to over-sample the minority class to create a balanced training set for model training.

In this study, we present a deep learning-based method, node2loc, to predict protein subcellular location. node2loc first learns distributed representation of proteins in a protein-protein network, which acquire effective representations from unlabeled data for downstream tasks. Then the learned representations are further fed into recurrent neural network (RNN) to classify 16 subcellular locations of proteins. Our results demonstrate that node2loc outperforms other baseline methods on our constructed benchmark dataset and yield promising performance.

## 2. Materials and Methods

We first construct a benchmark dataset with 16 subcellular locations. Then, node2vec is used to extract the embedding of proteins from the STRING network. In the end, the learned node embedding is further fed into a LSTM to classify 16 subcellular locations. The whole flowchart is shown in Figure 1.

**Figure 1.**
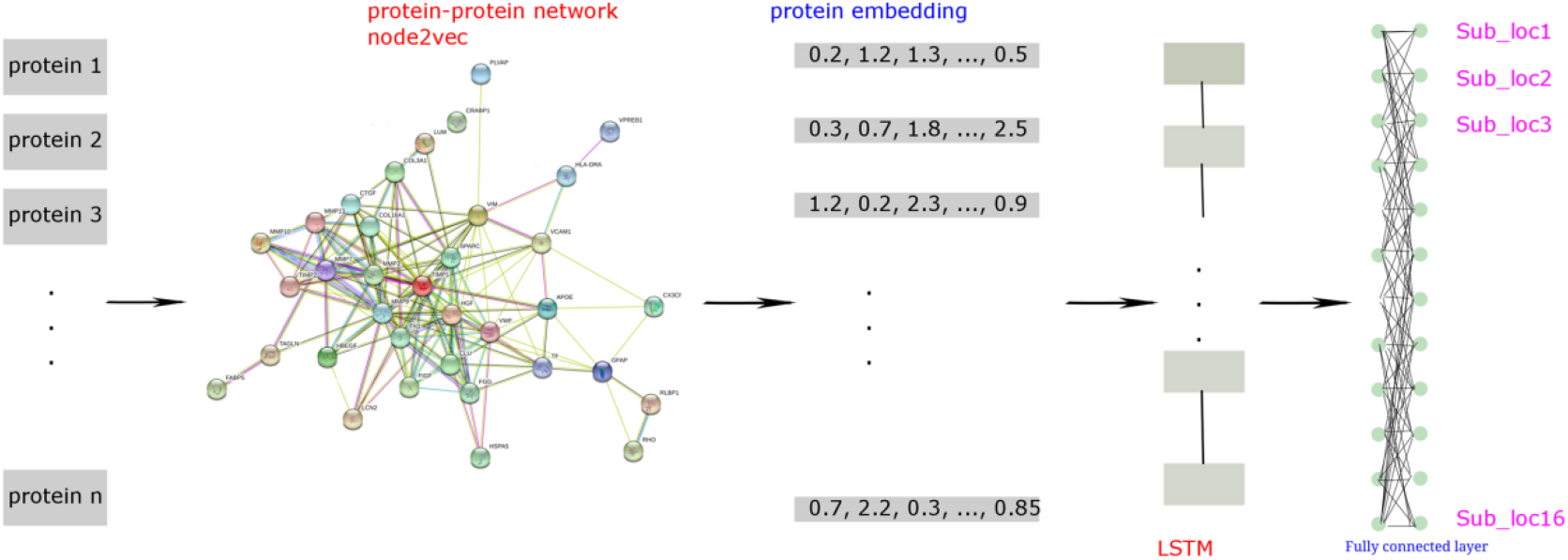
The flowchart of node2loc. node2loc first learns embedding of proteins from a protein-protein network using node2vec. Then the learned embedding is further fed into a RNN for classifying 16 subcellular locations. To reduce the impact of data imbalance on model training, we apply SMOTE to create a balanced training set. The network image is downloaded from STRING database.

### 2.1 Data sources

We employed the 5,960 protein sequences, represented by Uniprot IDs, used in one previous study [7], which were retrieved from Swiss-Prot (http://cn.expasy.org/, release 54.0) and refined by excluding proteins containing less than 50 amino acids or more than 5000 amino acids or unknown amino acids. Furthermore, these 5,960 protein sequences were with similarity less than 0.7 processed by CD-HIT [22]. Because we adopted a network embedding method to extract informative features of proteins from a protein-protein interaction (PPI) network reported in STRING [23], in which proteins were represented by Ensembl IDs, 5,960 Uniprot IDs were mapped onto their Ensembl IDs and those not occurring in the PPI network were discarded. Finally, 5,497 proteins were accessed. Their distribution on 16 categories is shown in Table 1. In addition, we also use CD-HIT to reduce the sequence similarity with a cutoff value 0.5. In total, we obtain 4,441 proteins and 16 subcellular locations. We obtain the same number of subcellular locations, but 1,056 fewer proteins than that of the similarity cutoff 0.7. The details of proteins and subcellular locations for similarity cutoff 0.5 are given in Table S1.

### 2.2 node2vec

In network biology, how to represent a node in the graph is an important topic. node2vec is developed to learn the continuous representation of nodes in a graph [17], then the learned representations can be used for downstream classification Word2vec [19] is used to learn the embedding of words by learning the co-occurrence of words in the training documents consisting of sequence of words. Thus, it first needs sampling a sequence of neighbor nodes from the graph. To efficiently explore the diverse neighborhoods, node2vec defines a flexible network neighborhood and designs a biased random walk approach.

To extend the Skip-gram of word2vec to network, node2vec aims to optimize the following objective function to maximize the log-probability of neighbor nodes given the new feature representation:

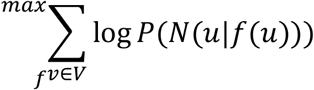

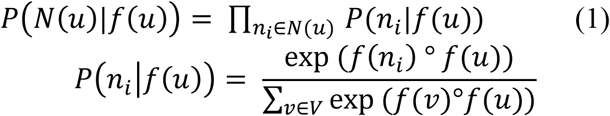

where N(u) is the neighborhood of node u, f is the new feature representation.

To solve the above objectives, node neighborhood need be first obtained. node2vec designs a flexible neighborhood sampling approach to improve the strategy of normal random walk by making tradeoff between the breadth-first sampling and depth-first sampling.

Given a source node s, a random walk of a fixed length is simulated according to below transition probability *π_uv_*:

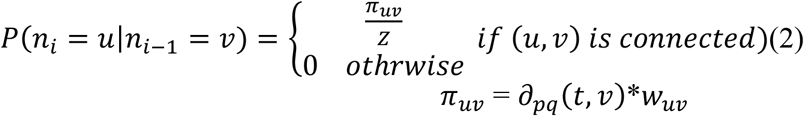

where *w_uv_* is the weight between two nodes.

node2vec defines a novel transition probability with return parameter p and in-out parameter q:

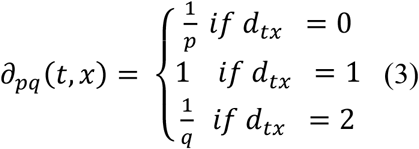

where *d_tx_* is the shortest path distance between nodes t and x.

After obtaining the node neighborhood, node2vec generates multiple sequences of nodes with a fixed length. These sequences of nodes are treated as sentences, and nodes as words. Then the widely used word2vec is ran on these pseudo sentences to generate the vector representations of nodes.

In this study, we used node2vec to learn embedding of proteins from a protein-protein network, which is collected from STRING (version 9.1). In total, the network has 2,425,314 interaction pairs from 20,770human proteins. We configure the dimension of embedding to 500, and other parameters are default values. The implementation of node2vec is downloaded from https://snap.stanford.edu/node2vec/.

### 2.3 Recurrent neural network

Recurrent neural network (RNN) is a type of neural network with loop inside for sequential data, whose output depends on the previous computations. The RNN allows information to persist for subsequential outputs, but in practice they are limited to looking back only a few steps.

Long short term memory network (LSTM) [13] is a type of RNN, it is capable of learning long-term dependencies. There exist three types of layers in LSTM, they are forget gate layer, input gate layer and output layer. The forget gate layer decides which information of previous state should be ignored. The input gate layer is used to determine which information should be passed to subsequent layer. The output gate layer decides what parts of state value should be outputted. In below text, we use the RNN to refer to the LSTM.

### 2.4 Reorder the learned embedding

In this study, we learn the node embedding with 500 dimensions from a protein-protein network. However, there exists some redundancy in the 500-dimensional embedding for supervised classifiers, and not all 500-D embedding should be considered to have the same discriminate power and used for classifying subcellular locations, some dimensions are more important than others. Thus, we first apply Minimum redundancy maximum relevance (mRMR) method, to rank the 500-D embedding, then a series of feature subsets are constructed on the ranked feature list in the incremental feature selection (IFS) method[24].

Each feature subset has one more feature than the preceding feature subset. For each feature subset, one supervised classifier is trained on the samples consisting of the features from the feature subset, and evaluated using 10-fold cross-validation. In the end, the classifier with the best performance is used to report the final performance and its corresponding feature subset is used.

### 2.5 Synthetic Minority Over-sampling Technique

As listed in Table 1, some categories (e.g., “Biological membrane”) contain lots of proteins, while some categories (e.g. “Flagellum or cilium”) have few samples, which induce difficulties for constructing effective prediction models. Here, we adopt the SMOTE [20] to tackle this problem. SMOTE is an over-sampling method, which can produce predefined number of new samples from the samples in a category with small size. The producing procedure can be described as follows. For each sample x in the category, calculate its Euclid distance to all other samples in this category. Then, select k (k is a parameter) nearest neighbors of this sample. Randomly choose one neighbor from these neighbors, say y, to produce a new sample z by the following equations:

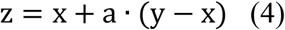

where *a* is a random number between 0 and 1. Because the new sample is generated by two similar samples in the same category, the new sample is also put into this category. Weka is a suite of software collecting several machine learning algorithms and data processing tools[25]. The tool, named “SMOTE”, implements the SMOTE method mentioned above. In this study, it is directly used for convenience. In detail, for each category expect “Biological membrane”, we generate some new samples via “SMOTE” and put them into this category. To construct a balanced dataset, each category contained almost same number of samples by adding new pseudo samples generated by “SMOTE”.

### 2.6 Baseline methods

To demonstrate the advantage of node2loc, we also make another baseline method STRING-KNN, which directly make use of confidence score from STRING database. For a given protein, 1) STRING-KNN finds the K interacting proteins from the training set with K largest confidence scores from its all interacting proteins;2) For each of 16 subcellular locations, based on the K proteins, STRING-KNN calculates the mean scores of confidence scores from the proteins with this subcellular location. If there is no interacting protein with this location, then the score is 0. We obtain 16 scores for this protein;3) We uses the subcellular location corresponding to the largest score as the predicted location for this protein. In this study, we try the K values 1,2,3,4,5,6,7,8,9,10,15,20,25,30,35,40,45 and 50.

In this study, we use RNN as the classifier of node2loc, we also evaluate other two widely used classifiers: random forest (RF)[26] and support vector machine (SVM)[27], which both use the learned embedding of proteins as inputs. For SVM, we test two different kernel functions:PolyKernel and Radial Basis Function (RBF),and the default values of other parameters in Weka are used. ForRF, we test three values 50, 100, 150 for parameter number of trees, and other parameters are default values in Weka. For RNN, we use epochs 500 and hidden number of neurons 400.

### 2.7 Performance measurement

In this study, we use 10-fold cross-validation to evaluate the performance of trained multiclass classifiers. We calculate the accuracy for individual class and overall accuracy for all classes. To be more objective, we also calculate the Matthews correlation coefficient (MCC)[28], a generalized version of original MCC that can only deal with the predicted results of binary classification. Assume we have *N* samples (*i* = 1, 2, …, *N*) and *C* classes (*j* = 1, 2, …, *C*), *X* = (*x_ij_*)_*N×C*_ is a matrix for the predicted classes of samples, where each *x_ij_* is 0 or 1; *x_ij_* equals to 1 if the sample *i* is predicted to be class *j*; otherwise 0. The matrix *Y* = (*y_ij_*)_*N×C*_ is the matrix for the true classes of samples, where the binary variable *y_ij_* = 1 when the sample *i* belongs to class *j*; otherwise 0. The MCC is defined as:

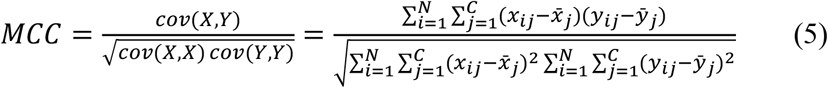

where 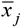 and 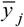 are the mean values of numbers of *x_j_* and *y_j_*, respectively.

## 3. Results

In this section, we first visualize the learned embedding of proteins from the protein-protein interaction network. Then we evaluate the performance of node2loc for classifying the 16 subcellular locations and compare node2loc with other baseline methods.

### 3.1 The learned node embedding of proteins

We first investigate the learned representations of proteins from a protein-protein network. Here we use t-SNE [29] to reduce the dimension of protein embedding from 500 to 2, which is further visualized in the 2-D space. The label information for proteins are from our benchmark dataset. As shown in Figure 2A, the learned embedding is to some extent divided into different groups, some clusters correspond to subcellular locations. For example, the embedding of proteins with “biological membrane” is clearly different from proteins from other locations. In addition, we also illustrate the K-means clustering results of embedding of 22,070 proteins (Figure 2B). The results indicate that the low dimensional embedding learned by node2vec still have certain discriminate ability for classifying different subcellular locations, even the embedding is learned in a completely unsupervised way.

**Figure 2.**
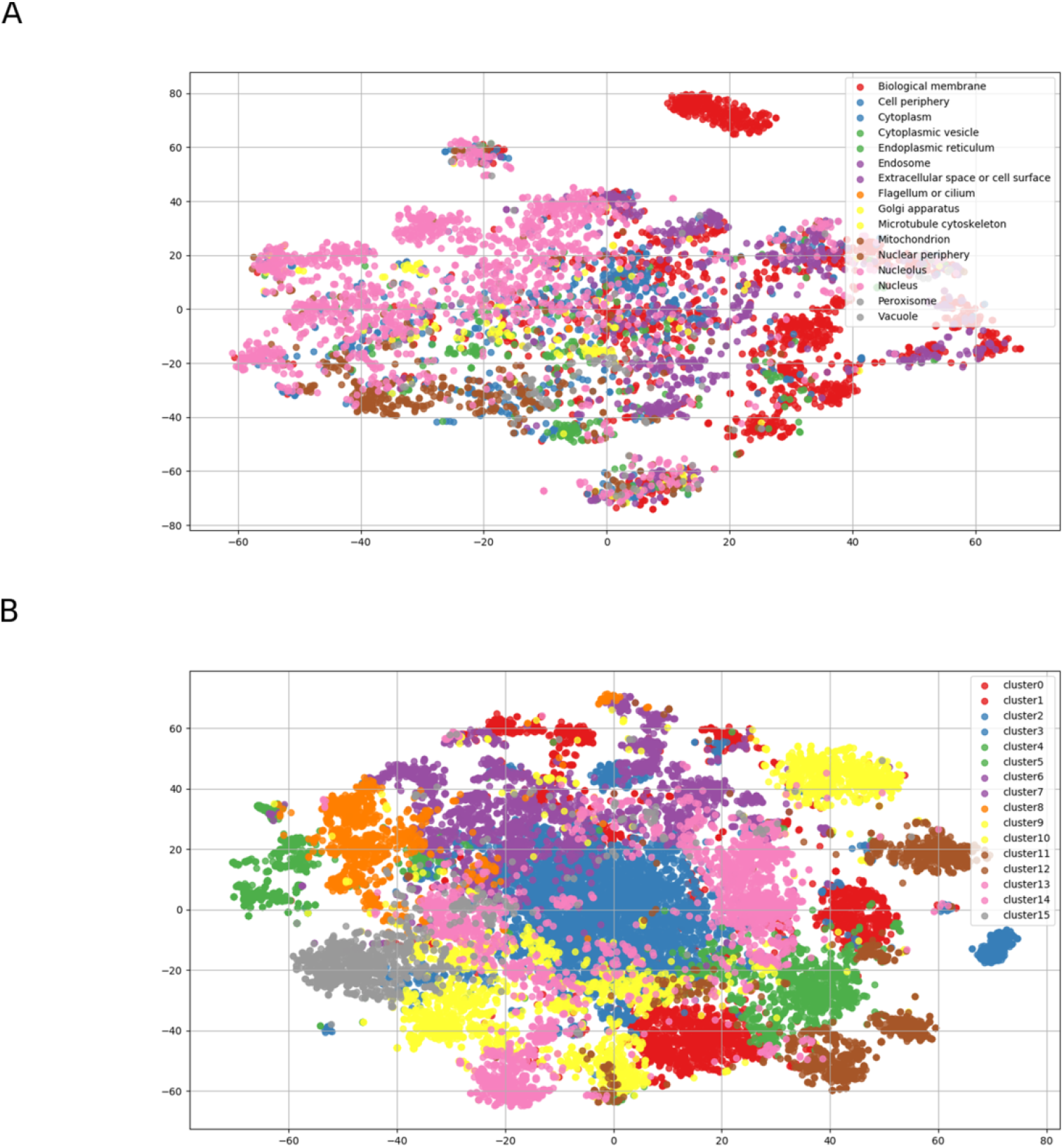
Visualization of learned embedding in 2-D space using t-SNE. A) The learned embedding of proteins in our benchmark set, the labels are subcellular locations of proteins from the benchmark dataset. B) The learned embedding of all 20,770 proteins, the labels are the K-means clustering results with number of clusters K=16.

### 3.2 The performance of node2loc

We classify the 16 subcellular locations by combining the learned embedding of proteins and an RNN. In this study, we learned 500-dimensional embedding from the network. We use IFS method with the RNN to evaluate the performance using feature subsets with the number of features from 1 to 500 according to the feature list generated by mRMR, where feature subset 1 has the top 1 feature, and feature subset 2 has top 1 and 2 features, and so on. SMOTE is used to reduce the impact of imbalanced classes. As shown in Figure 3, node2loc achieves a best MCC value of 0.812 when using top 497-D embedding. And node2loc achieves an MCC value of 0.80 when using the whole 500-D embedding. In addition, node2loc yields an overall accuracy of 0.843 across the 16 subcellular locations. Compared to our previous study [7] that uses features derived from functional data, the overall accuracy of node2loc is much higher than 0.677 in our previous study [7], it is an increase by 24.5%.

**Figure 3.**
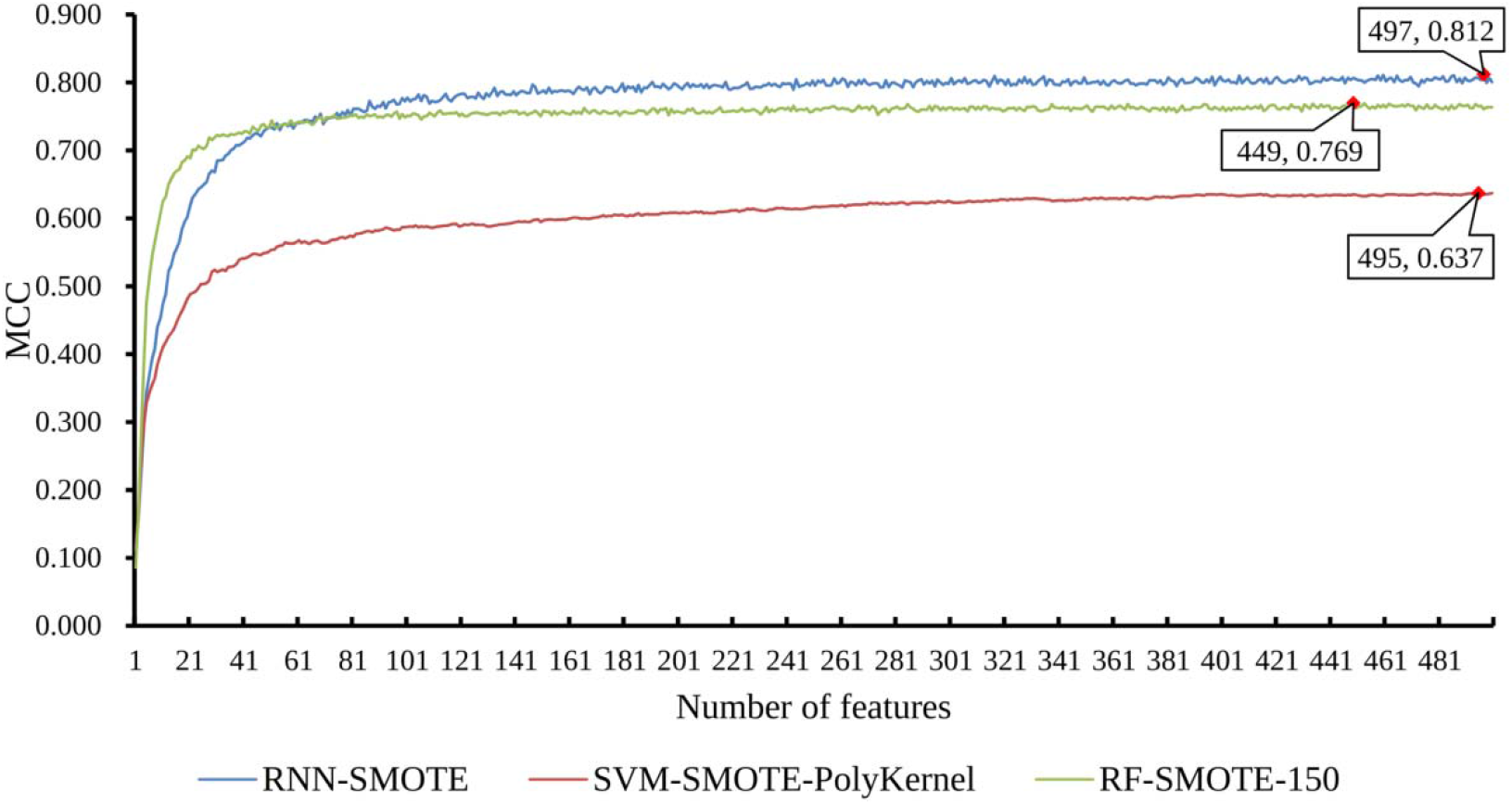
The classification performance of different classifiers RNN, RF and SVM when using different dimensions of learned embedding and SMOTE. RNN-SMOTE is our method node2loc, SVM-SMOTE-PolyKernel means the SVM uses PolyKernel. RF-SMOTE-150 means RF uses number of trees 150.

In this study, node2loc uses RNN as the supervised classifier. To demonstrate the advantage of the RNN, we also used other two classifiers: SVM [27] and RF [26], which also use the learned embedding of proteins as inputs and SMOTE for over-sampling. As shown in Table 2, SVM yields better performance using PolyKernal than RBF kernel. And RF yields better performance using number of trees 150 than 50 and 100, respectively. Figure S1 and Figure S2 illustrate the change of the MCC with different dimensions of learned embedding when using different parameters for RF and SVM, respectively.

As shown in Figure 3, the RNN outperforms SVM and RF. RF yields an MCC value of 0.769 when using 449-D embedding, and SVM yields a much lower MCC value 0.637 when using 495-Dembedding. Both RF and SVM perform worse than the MCC 0.812 of RNN (Table 2). As shown Table S2, of the 16 subcellular locations, node2loc outperforms RF and SVM on almost all 16 locations (expect the class “Extracellular space or cell surface”) (Figure 4). The results indicate that RNN can be a more powerful classifier than conventional machine learning classifiers SVM and RF.

**Figure 4.**
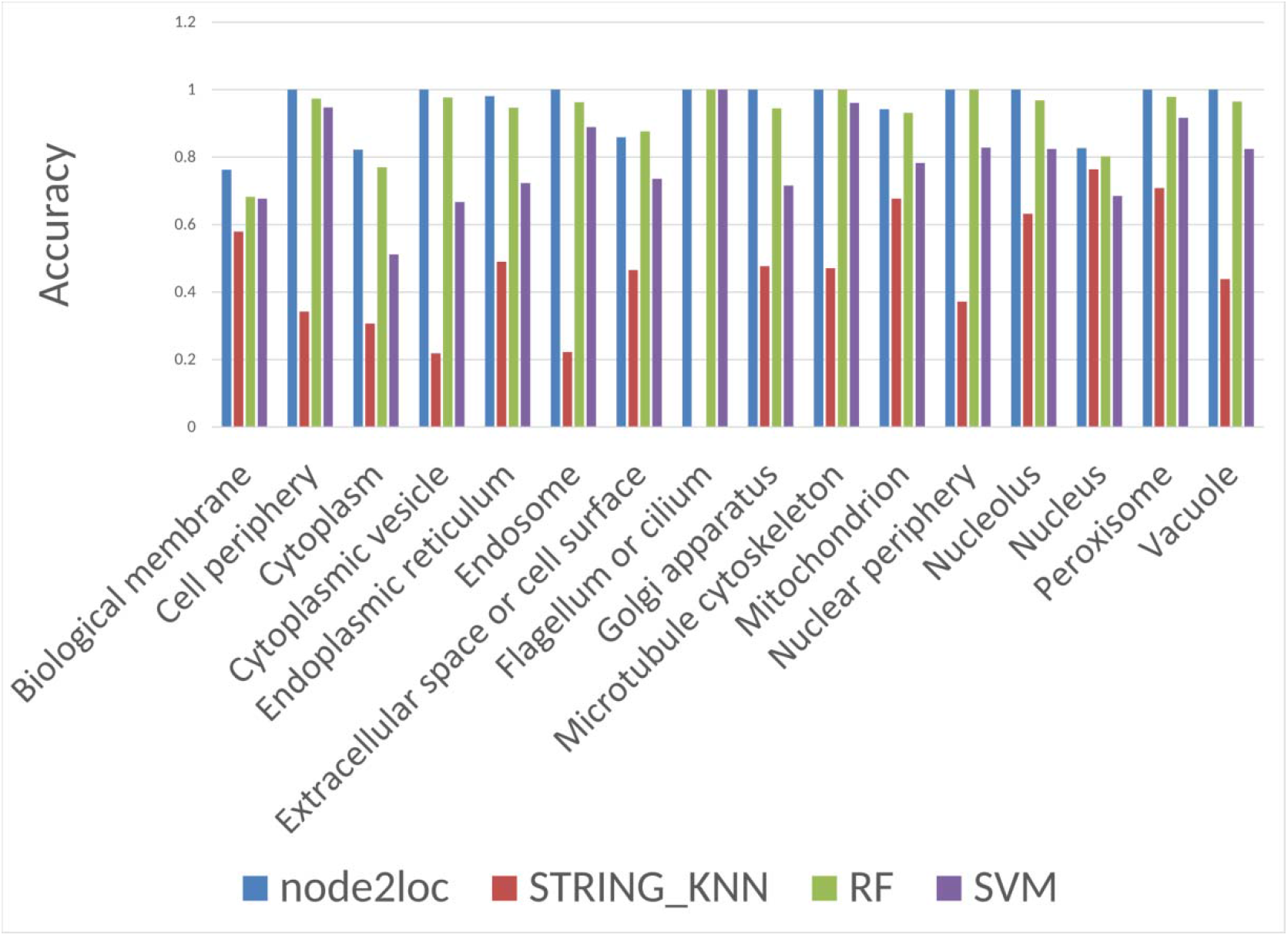
Performance comparison of different computational methods for 16 locations. node2loc, RF, SVM use learned embedding as inputs and SMOTE for oversampling.

In addition, we evaluate our methods on a dataset with a CD-HIT removal of sequence similarity cutoff 0.5. node2loc achieves an overall accuracy of 0.816 and an MCC value of 0.783 when using 462-dimensional embedding and the performanceis a little lower than the performance on our benchmark set with a cutoff 0.7. Figure S3 shows the performance MCCs change with the dimension of embedding. The results demonstrate that the sequence similarity has no big impact on the performance of node2loc. It should be noted that node2loc does not use any sequence-derived features for model training, it only uses the learned embedding from a protein-protein network for classifying subcellular locations.

In this study, we learn the node embedding from a human protein-protein network with 20,770human proteins from STRING database. We also learn the node embedding from a small protein-protein network, in which only those interactions among the 5,497 proteins in the benchmark dataset are kept. When using learned node embedding from this small protein-protein network, node2loc yields an MCC value of 0.782, which is lower than an MCC value of 0.812 using node embedding from the large human STRING protein-protein network. The results indicate that node2vec can learn better node embedding from a larger protein-protein network.

### 3.3 Comparing node2loc with another baseline method STRING-KNN

We first evaluate the impact of the parameter K on the prediction performance for STRING-KNN. The performance using different K values are given in Table S3, and STRING-KNN yields the best MCC value of 0.476 when K=1. As shown in Table 2, STRING-KNN achieves an MCC value of 0.476, which is much lower than the MCC0.812 of node2loc. STRING-KNN yields the overall accuracy of 0.575, which is also much lower than 0.843 of node2loc. node2loc outperforms STRING-KNN by an increase of over 50%. We also check the prediction accuracy of STRING-KNN on 16 individual locations. As shown in Figure 4 and Table S2, node2loc outperforms STRING-KNN on all 16 subcellular locations. Especially for those minority classes “Flagellum or cilium”, node2loc can classify samples of this location with 100% accuracy; however, STRING-KNN completely cannot classify the samples of this location. The results indicate that node2loc yields very promising performance on predicting 16 subcellular locations, and performs much better than the baseline method STRING-KNN.

### 3.4 The added value of over-sampling SMOTE

In node2loc, we applied SMOTE to reduce the impact of data imbalance. As shown in Figure 5, over-sampling using SMOTE can improve the prediction performance, especially for those minority classes. For node2loc, SMOTE improves the MCC from 0.615 to 0.812, which is an increase by 32.0%. For RF, SMOTE also increases the performance from MCC 0.616 to 0.769. SMOTE improves the overall performance of SVM with a much smaller margin, it increases the MCC value from 0.623 to 0.637. The MCC changes with the different dimensions of learned embedding as inputs when using SMOTE or not for RF (Figure S1), SVM (Figure S2) and node2loc (Figure S4), respectively. For RF and node2loc, SMOTE always improves the prediction performance. The results show that the over-sampling samples in embedding space can make the trained models be able to classify those samples from minority classes. It is because that learned embedding takes interacting proteins into account, and interacting proteins tends to be close in embedding space. In addition, interacting proteins tend to locate in the same subcellular location, thus over-sampling methods can introduce better synthesized samples.

**Figure 5.**
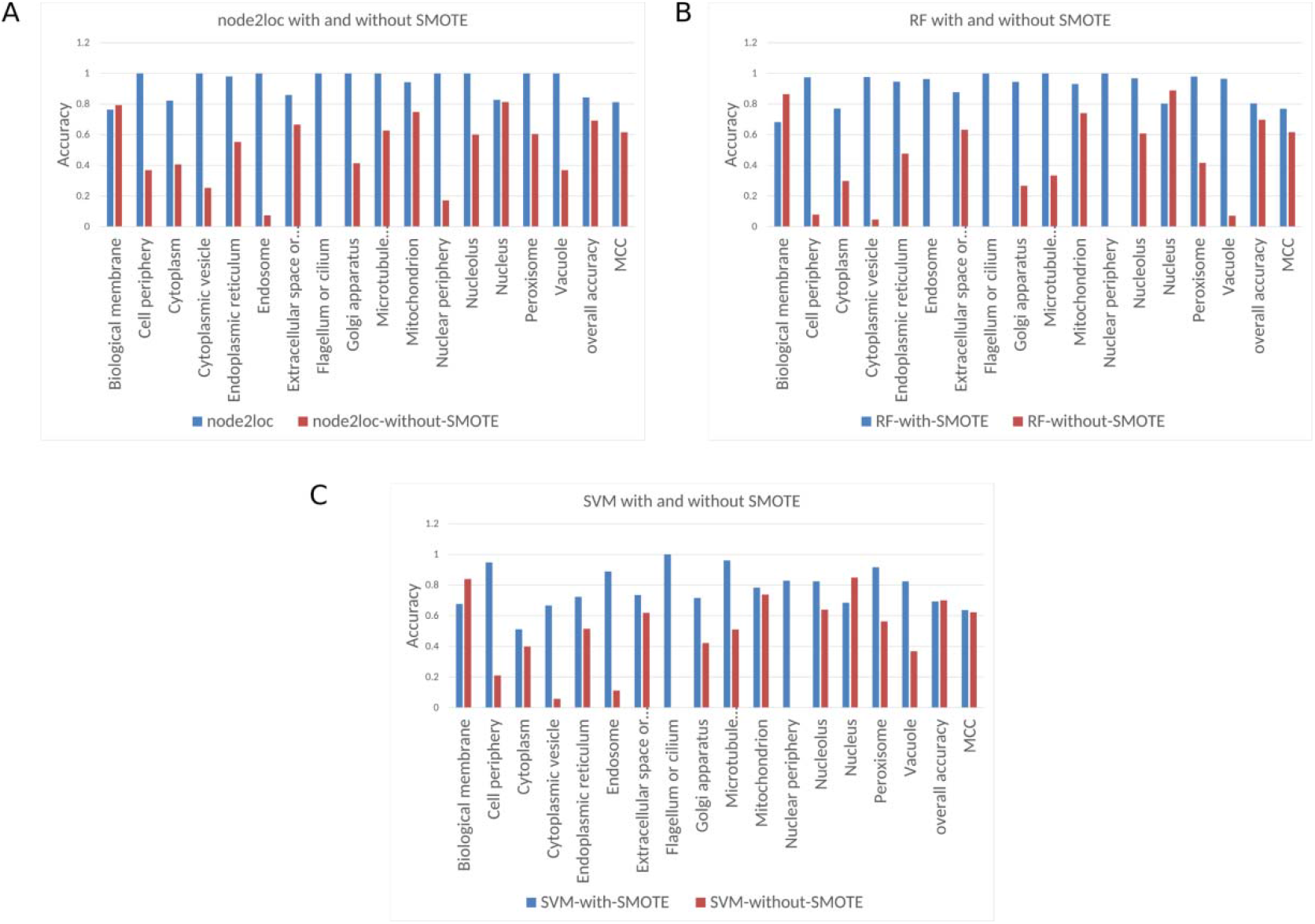
The performance comparison of node2loc, RF, SVM using SMOTE or not. A) Accuracy of individual locations, overall accuracy and MCC for node2loc when using SMOTE and not using SMOTE; B) Accuracy of individual locations, overall accuracy and MCC for RF when using SMOTE and not using SMOTE; C) Accuracy of individual locations, overall accuracy and MCC for SVM when using SMOTE and not using SMOTE.

As shown in Table S4 and Figure 5, for those subcellular locations with small number of proteins, node2loc-without-SMOTE yields a very low accuracy. For example, node2loc-without-SMOTE yields an accuracy of 0.074 for “Endosome” with 27 proteins and 0 for “Flagellum or cilium” with three proteins. After over-sampling the minority class using SMOTE, node2loc improves the accuracy to 1 for both “Endosome” and “Flagellum or cilium”. Over-sampling using SMOTE improves the prediction accuracy of node2loc for 15 locations, and it only reduces prediction accuracy for the majority location “Biological membrane” by a small margin, which has over 1000 proteins. In addition, the overall accuracy of node2loc is 0.843, which is much higher than 0.692 of node2loc-without-SMOTE. SVM and RF have the similar results. The results indicate that SMOTE hugely reduce the impact of data imbalance and improve the prediction performance of node2loc by a large margin.

### 3.5 Proteome-wide subcellular locations prediction

In this study, we learned node embedding for 20,770 proteins, of them, 5,497 proteins with annotated subcellular locations are used for model training, the 15,273 proteins are left. Of the 15,273 proteins, 100 proteins have sequence similarity > 70% to the benchmark set, and they have annotated locations but are removed from model training. We also filter out these 100 proteins from the remaining 15,273 proteins. Thus, we obtain 15,173 proteins that have no annotated subcellular locations. We use node2loc trained on 5,497 proteins to predict subcellular locations for these 15,173 proteins. The predicted subcellular locations for them using node2loc are given in Table S5. As shown in Figure 6, node2loc can predict more proteins with the minority subcellular locations in the benchmark set than node2loc without SMOTE. In addition, node2loc without SMOTE is preferred to predict proteins with the majority subcellular locations. The results also demonstrate that SMOTE can make the trained model be not biased to the majority classes.

**Figure 6.**
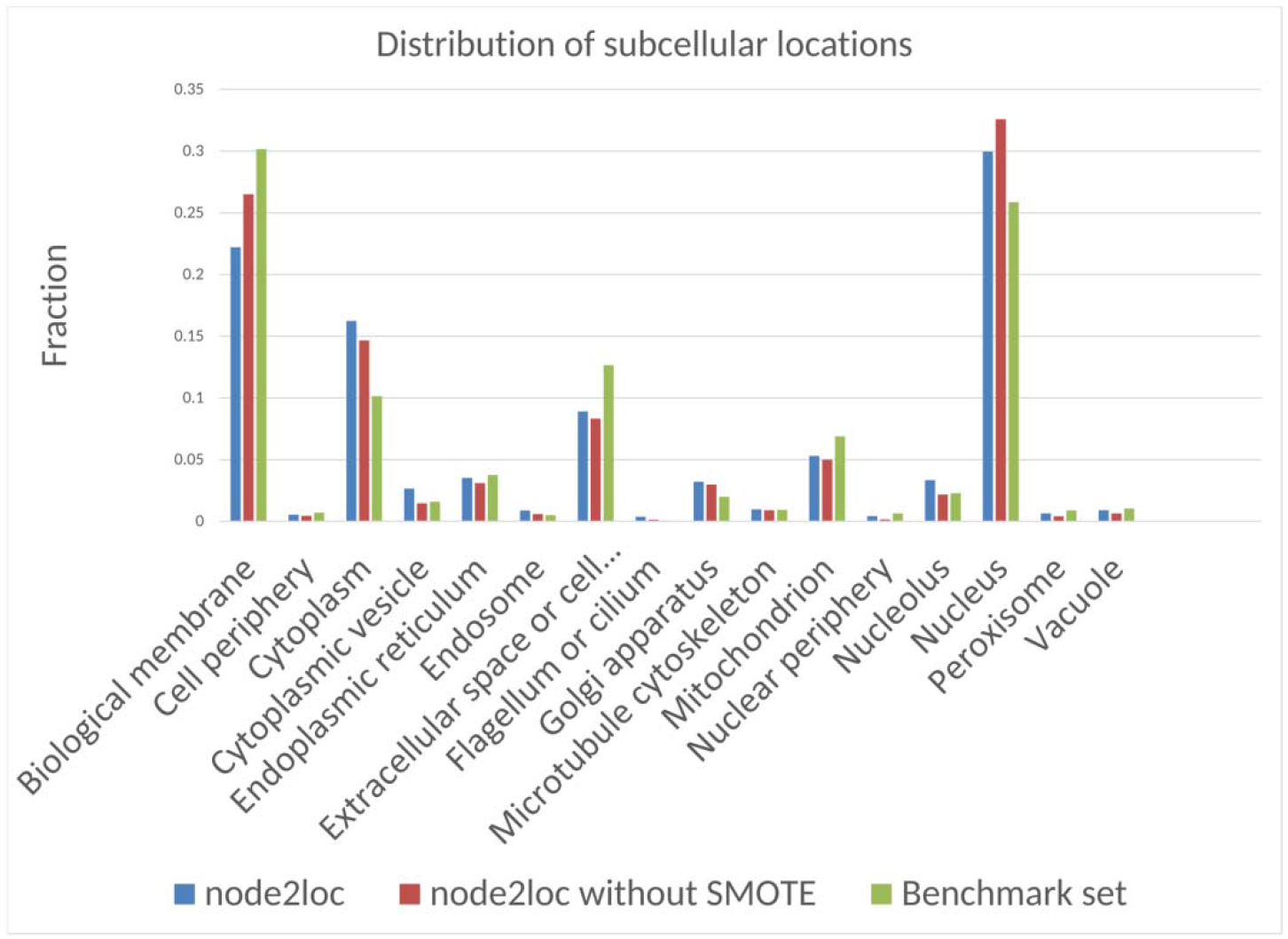
The comparison of fractions of subcellular locations predicted by node2loc, node2loc without SMOTE and the benchmark set.

We further investigate predicted subcellular locations of some proteins by node2loc, and these proteins are not in our benchmark set. We first search the protein in COMPARTMENTS database [30], which gives the evidence of locations from multiple channels, including knowledge, experiments and text mining. As shown in Table 3, we search locations for some proteins in COMPARTMENTS and find the support evidence for the predicted locations by node2loc (Table 3). For each of the 16 locations, there are proteins predicted to be located in it accurately by node2loc. The results indicate that node2loc can make accurate subcellular localization predictions for some unseen proteins in the benchmark set.

## 4. Discussion

Our results demonstrated that the learned embedding from a network have strong discriminatory power for downstream classification. When learning the embedding from the protein-protein network, we do not take any information related to subcellular location into account. In addition, the learned embedding is learned in a completely unsupervised way, it can also be applied for other downstream tasks, e.g. proteins’ functions prediction.

The learned node embedding are still black-box and we cannot trace it back to which features are important for classifying subcellular locations. For subcellular location prediction, we pay more attention to the prediction accuracy. But if we can have more details about why the model makes such a decision, then the prediction will be more convincing.

One disadvantage of node2loc is that it requires the proteins have known interacting proteins in STRING database. For those proteins without binding proteins, we discard 463 proteins from our training samples (section 2.1) since we cannot learn node embedding for them. On the other hand, we need retrain the node2vec on the whole protein-protein network with unseen proteins if we want to predict locations for new proteins. Recently, a new inductive method GraphSAGE[18] can learn functions to compute node representations for unseen nodes. In future work, we expect to use GraphSAGE for learning embedding for proteins.

In this study, we demonstrate that over-sampling samples using SMOTE in embedding space can improve the prediction performance with a large margin, especially for those minority classes. Instead of duplicating samples from minority classes and under-sample the majority classes, SMOTE generate synthetic samples in “feature space” rather than “data space”. SMOTE forces the decision boundary of minority classto be more general[20].SMOTE makes the trained models to be not biased to the majority classes or the minority classes, as shown in Figure 6. There exist other over-sampling methods, e.g. Supervised Over-Sampling [21], which shows better performance than SMOTE. In future work, we expect to improve the prediction performance by: 1) developing more advanced methods to reduce the impact of data imbalance. 2) using data augmentation, which is widely used in deep learning-based image processing. 3) exploring the alignment among two or multiple interaction graphs [31, 32] from different species using graph embedding, instead of using only human interaction network. 4) using an end-end learning framework graph convolutional network [33] to directly infer the protein subcellular locations in the protein-protein network. 5) node2loc considers each protein has only one location, however, some proteins have multiple locations. We can format the task as multi-label learning, which can predict multiple locations for one protein. node2loc can be easily adapted for multi-label classification, we just need modify the activation function of the last layer to be sigmoid, and the loss function uses binary cross entropy.

## 5. Conclusion

In this study, we present a deep learning-based method to predict protein subcellular location under the context of protein-protein network. Our method node2loc directly learns node embedding from protein-protein network in an unsupervised way. Then these learned embedding are fed into supervised classifiers to classify 16 subcellular locations. To reduce the impact of data imbalance, we use the over-sampling method SMOTE to create balanced training set in embedding space to improve the prediction performance. Our results demonstrate that the learned embedding is effective for downstream subcellular location classification. Combining with an RNN, node2loc outperforms other three baseline methods on our constructed benchmark set. In addition, we do a large-scale subcellular location prediction for those proteins not in the benchmark set, and some predictions are supported by literature and public databases.

## Acknowledgements

This study was supported by the the Shanghai Municipal Science and Technology Major Project (2017SHZDZX01), National Natural Science Foundation of China [31701151, 61903248], Natural Science Foundation of Shanghai [17ZR1412500], National Key R&D Program of China [2018YFC0910403], Shanghai Sailing Program [16YF1413800], the Youth Innovation Promotion Association of Chinese Academy of Sciences (CAS) [2016245], the fund of the key Laboratory of Stem Cell Biology of Chinese Academy of Sciences [201703], Science and Technology Commission of Shanghai Municipality (STCSM) [18dz2271000]

## Reference

[1] K. Bimpikis, A. Budd, R. Linding et al., “BLAST2SRS, a web server for flexible retrieval of related protein sequences in the SWISS-PROT and SPTrEMBL databases,” Nucleic Acids Res, vol. 31, no. 13, pp. 3792–4, Jul 1, 2003.

[2] K. C. Chou, and H. B. Shen, “Euk-mPLoc: a fusion classifier for large-scale eukaryotic protein subcellular location prediction by incorporating multiple sites,” J Proteome Res, vol. 6, no. 5, pp. 1728–34, May, 2007.

[3] K.-J. Park, and M. Kanehisa, “Prediction of protein subcellular locations by support vector machines using compositions of amino acids and amino acid pairs,” Bioinformatics, vol. 19, no. 13, pp. 1656–1663, 2003.

[4] K. C. Chou, and Y. D. Cai, “Using functional domain composition and support vector machines for prediction of protein subcellular location,” J Biol Chem, vol. 277, no. 48, pp. 45765–9, Nov 29, 2002.

[5] K. C. Chou, and Y. D. Cai, “Prediction and classification of protein subcellular location-sequence-order effect and pseudo amino acid composition,” J Cell Biochem, vol. 90, no. 6, pp. 1250–60, Dec 15, 2003.

[6] H. Zhou, Y. Yang, and H. B. Shen, “Hum-mPLoc 3.0: prediction enhancement of human protein subcellular localization through modeling the hidden correlations of gene ontology and functional domain features,” Bioinformatics, vol. 33, no. 6, pp. 843–853, Mar 15, 2017.

[7] B.-Q. Li, T. Huang, L. Chen et al., “Prediction of Human Protein Subcellular Locations with Feature Selection and Analysis,” Frontiers in Protein and Peptide Sciences, B. M. Dunn, ed., pp. 206–225, Soest: Bentham Science Publishers, 2014.

[8] J. Q. Jiang, and M. Wu, “Predicting multiplex subcellular localization of proteins using protein-protein interaction network: a comparative study,” BMC Bioinformatics, vol. 13 Suppl 10, pp. S20, Jun 25, 2012.

[9] B. Alipanahi, A. Delong, M. T. Weirauch et al., “Predicting the sequence specificities of DNA-and RNA-binding proteins by deep learning,” Nat Biotechnol, vol. 33, no. 8, pp. 831–8, Aug, 2015.

[10] Y. Lecun, L. Bottou, Y. Bengio et al., “Gradient-based learning applied to document recognition,” Proceedings of the Ieee, vol. 86, no. 11, pp. 2278–2324, Nov, 1998.

[11] X. Y. Pan, and H. B. Shen, “RNA-protein binding motifs mining with a new hybrid deep learning based cross-domain knowledge integration approach,” Bmc Bioinformatics, vol. 18, pp. 136, Feb 28, 2017.

[12] X. Pan, P. Rijnbeek, J. Yan et al., “Prediction of RNA-protein sequence and structure binding preferences using deep convolutional and recurrent neural networks,” BMC Genomics, vol. 19, pp. 511, 2018.

[13] S. Hochreiter, and J. Schmidhuber, “Long short-term memory,” Neural computation, vol. 9, no. 8, pp. 1735–1780, 1997.

[14] X. Pan, and H. B. Shen, “Predicting RNA-protein binding sites and motifs through combining local and global deep convolutional neural networks,” Bioinformatics, vol. 34, no. 20, pp. 3427–3436, Oct 15, 2018.

[15] J. Zhou, and O. G. Troyanskaya, “Predicting effects of noncoding variants with deep learning-based sequence model,” Nat Methods, vol. 12, no. 10, pp. 931–4, Oct, 2015.

[16] D. Quang, and X. Xie, “DanQ: a hybrid convolutional and recurrent deep neural network for quantifying the function of DNA sequences,” Nucleic Acids Res, vol. 44, no. 11, pp. e107, Jun 20, 2016.

[17] A. Grover, and J. Leskovec, “node2vec: Scalable Feature Learning for Networks,” in Proceedings of the 22nd ACM SIGKDD International Conference on Knowledge Discovery and Data Mining, San Francisco, California, USA, 2016, pp. 855–864.

[18] W. Hamilton, Z. Ying, and J. Leskovec, “Inductive representation learning on large graphs.” pp. 1024–1034.

[19] T. Mikolov, I. Sutskever, K. Chen et al., “Distributed representations of words and phrases and their compositionality.” pp. 3111–3119.

[20] N. V. Chawla, K. W. Bowyer, L. O. Hall et al., “SMOTE: Synthetic minority oversampling technique,” Journal of Artificial Intelligence Research, vol. 16, pp. 321–357, 2002.

[21] Z. Cao, X. Pan, Y. Yang et al., “The lncLocator: a subcellular localization predictor for long non-coding RNAs based on a stacked ensemble classifier,” Bioinformatics, vol. 34, no. 13, pp. 2185–2194, Jul 1, 2018.

[22] W. Li, and A. Godzik, “Cd-hit: a fast program for clustering and comparing large sets of protein or nucleotide sequences,” Bioinformatics, vol. 22, no. 13, pp. 1658–9, Jul 1, 2006.

[23] D. Szklarczyk, A. Franceschini, S. Wyder et al., “STRING v10: protein-protein interaction networks, integrated over the tree of life,” Nucleic Acids Res, vol. 43, no. Database issue, pp. D447–D452, Oct 28, 2015.

[24] H. A. Liu, and R. Setiono, “Incremental feature selection,” Applied Intelligence, vol. 9, no. 3, pp. 217–230, Nov-Dec, 1998.

[25] Author ed.^eds., “Data Mining:Practical Machine Learning Tools and Techniques,” 2nd edn, San Francisco, Morgan, Kaufmann, 2005, p.^pp. Pages.

[26] L. Breiman, “Random forests,” Machine learning, vol. 45, no. 1, pp. 5–32, 2001.

[27] C. Cortes, and V. Vapnik, “Support-vector networks,” Machine Learning, vol. 20, no. 3, pp. 273–297, 1995.

[28] J. Gorodkin, “Comparing two K-category assignments by a K-category correlation coefficient,” Computational Biology and Chemistry, vol. 28, no. 5, pp. 367–374, 2004.

[29] M. LJPvd, and G. Hinton, “Visualizing high-dimensional data using t-SNE,” Journal of Machine Learning Research, vol. 9, pp. 2579–605, 2008.

[30] J. X. Binder, S. Pletscher-Frankild, K. Tsafou et al., “COMPARTMENTS: unification and visualization of protein subcellular localization evidence,” Database (Oxford), vol. 2014, pp. bau012, 2014.

[31] J. Yan, M. Cho, H. Zha et al., “Multi-graph matching via affinity optimization with graduated consistency regularization,” IEEE transactions on pattern analysis and machine intelligence, vol. 38, no. 6, pp. 1228–1242, 2016.

[32] J. Yan, J. Wang, H. Zha et al., “Consistency-driven alternating optimization for multigraph matching: A unified approach,” IEEE Transactions on Image Processing, vol. 24, no. 3, pp. 994–1009, 2015.

[33] T. N. Kipf, and M. Welling, “Semi-supervised classification with graph convolutional networks,” in 5th International Conference on Learning Representations, 2017.

